# SARS-CoV-2 Omicron Variant AI-based Primers

**DOI:** 10.1101/2022.01.21.475953

**Authors:** Carmina A. Perez-Romero, Alberto Tonda, Lucero Mendoza-Maldonado, John MacSharry, Joanna Szafran, Eric Claassen, Johan Garssen, Aletta D. Kraneveld, Alejandro Lopez-Rincon

**Affiliations:** Division of Pharmacology, Utrecht Institute for Pharmaceutical Sciences, Faculty of Science, Utrecht University, Universiteitsweg 99, 3584 CG Utrecht, the Netherlands; Departamento de Investigación, Universidad Central de Queretaro (UNICEQ), Av. 5 de Febrero 1602, San Pablo, 76130 Santiago de Querétaro, Qro., Mexico; UMR 518 MIA-Paris, INRAE, c/o 113 rue Nationale, 75103, Paris, France; Hospital Civil de Guadalajara “Dr. Juan I. Menchaca”. Salvador Quevedo y Zubieta 750, Independencia Oriente, C.P. 44340 Guadalajara, Jalisco, México; Athena Institute, Vrije Universiteit, De Boelelaan 1085, 1081 HV Amsterdam, the Netherlands; Department Immunology, Danone Nutricia research, Uppsalalaan 12, 3584 CT Utrecht, the Netherlands; Julius Center for Health Sciences and Primary Care, University Medical Center Utrecht, Universiteitsweg 100, 3584 CG Utrecht; School of Microbiology and School of Medicine, University College Cork, College Rd, University College, Cork, Ireland

## Abstract

As the COVID-19 pandemic continues to affect the world, a new variant of concern, B.1.1.529 (Omicron), has been recently identified by the World Health Organization. At the time of writing, there are still no available primer sets specific to the Omicron variant, and its identification is only possible by using multiple targets, checking for specific failures, amplifying the suspect samples, and sequencing the results. This procedure is considerably time-consuming, in a situation where time might be of the essence. In this paper we use an Artificial Intelligence (AI) technique to identify a candidate primer set for the Omicron variant. The technique, based on Evolutionary Algorithms (EAs), has been already exploited in the recent past to develop primers for the B.1.1.7/Alpha variant, that have later been successfully tested in the lab. Starting from available virus samples, the technique explores the space of all possible subsequences of viral RNA, evaluating them as candidate primers. The criteria used to establish the suitability of a sequence as primer includes its frequency of appearance in samples labeled as Omicron, its absence from samples labeled as other variants, a specific range of melting temperature, and its CG content. The resulting primer set has been validated in *silico* and proves successful in preliminary laboratory tests. Thus, these results prove further that our technique could be established as a working template for a quick response to the appearance of new SARS-CoV-2 variants.

## Introduction

On November 26th, 2021, the World Health Organization (WHO) declared a fifth variant of concern (VOC)^1^, the Omicron variant, belonging to the PANGO lineage as B.1.1.529^2^, and clade GR/484A in the Global Initiative on Sharing Avian Influenza Data (GISAID)^3^. The Omicron variant is characterized by the mutations^4,5^ presented in Table 1.

**Table 1.** Characteristic Mutations of Omicron Variant

| Gene | Mutations |
| --- | --- |
| S | A67V, Δ69-70, T95I, G142D, Δ143-145, Δ211, L212I, ins214EPE, G339D, S371L, S373P, N764K, D796Y, N856K, Q954H, N969K, L981F, S375F, K417N, N440K, G446S, S477N, T478K, E484A, Q493R, G496S, Q498R, N501Y, Y505H, T547K, D614G, H655Y, N679K, P681H, |
| ORF1ab | K38R, V1069I, Δ1265, L1266I, A1892T, T492I, P132H, Δ105-107, A189V, P323L, I42V |
| E | T9I |
| M | D3G, Q19E, A63T |
| N | P13L, Δ31-33, R203K, G204R |

B.1.1.529 was first discovered in Botswana on November 11th, 2021. It was then quickly identified in South Africa three days later and identified in two cases in Hong Kong^4^. B.1.1.529 is characterized by at least 30 amino acid substitutions, three small deletions and one small insertion in the viral RNA encoding for the spike protein (S), making it the most divergent variant detected to date^4^. Importantly, half of the amino acid substitutions in B.1.1.529 are present on its receptor-binding domain (RBD), playing a key role in ACE2 binding and antibody recognition. Several mutations in this domain have been characterized in other variants, including VOCs^6,7^ and modelled associating them with increased binding affinity to ACE2 and higher rates of transmission (N501Y, Q498R, H655Y, N679K, P681H)^8–11^, increase in the ability of immune escape from neutralising antibodies (K417N, N501Y, N440K, S477N, T478K, G339D)^11–14^, reduction in vaccine effectiveness and increased risk of reinfections (E484, K417N, N501Y)^13–17^. The Omicron Variant has been divided into 3 sublineages: BA.1, BA.2 and BA.3, where as of December 15th, 2021 BA.1 accounts for 95% of available Omicron sequences in GISAID. BA.2 does not present the 69-70 deletion in the S gene, while BA.1 and BA.3 do^18^.

Since its discovery, B.1.1.529 has been rapidly spreading across the globe and through communities pointing to a higher degree of transmission and possible growth advantage when compared to other variants such as Delta (B.1.617.2)^15,19^. Although a great deal remain to be uncovered regarding clinical presentation and epidemiology, the majority of cases reported to authorities are either asymptomatic at the time of testing or have mild symptoms (including cough, fatigue, and congestion or runny nose)^4,20,21^: preliminary data thus suggest a lower disease severity^22,23^. Since its detection, B.1.1.529 has started to replace the Delta variant in South Africa^24–26^, and early evidence has linked it to an increased risk of reinfection^15–17,27^. Due to its highly mutated S protein, B.1.1.529 has been shown to escape the majority of SARS-CoV-2 neutralizing antibodies and retrovirals^12,13,28^ (except for ensovibep^29^, remdesivir, molnupiravir and nirmatrelvir^30,31^). Furthermore, current vaccines technologies tested against B.1.1.529 seem to show lower efficacy^13,23,26,28,32,33^, however vaccine boosters (including heterologous booster) have been shown to help reducing immune escape^31,34–37^, and universal vaccines to help fight this variant have been proposed^38,39^. It is important to note that early data point to a reduced risk of hospital admission among Omicron infected individuals^23,27^, and a reduced risk of severe outcomes among Omicron re-infected individuals earlier infected by the Delta variant^25^. Interestingly, a recent study found an increase in Delta variant neutralization from individuals infected with Omicron^40^, which could result in a decreased ability of reinfection with future and current variants.

Due to the high concern of this variant, one of the best strategies to contain its spread is by properly measuring it. The current identification of variants is usually performed through PCR tests. As we recently showed in^41^, on barriers influencing vaccine development timelines in Covid-19 Vaccine R&D, fast and correct identification is of highest impact on Covid-19 vaccine development. In other words, the steepest barrier to innovation is “lack of knowledge concerning the pathogen target”. Consequently, AI fast prediction of suitable primers relates directly to outbreak responses, as also recently discussed in^41,42^. Differentiating between Omicron and the other variants, however, is a difficult task. In practice, the main way of detecting the Omicron variant is by running three targets, and then checking the so-called S gene dropout or S gene target failure^1^: a failure of one specific test that targets the heavily mutated S gene. The virus is then amplified and sequenced, using a procedure similar to the one followed for identifying the B.1.1.7/Alpha variant^43^. Depending on the technology adopted and the laboratory conditions in each country, the whole process can take days. Considering the amount of the people that need to be tested in order to properly assess the spread of the variant, this creates the necessity for a faster and reliable test. In addition, if 69-70 deletion is targeted in the S gene, the BA.2 sublineage does not have this mutation. Here we present the design of a primer set for the specific detection of the Omicron variant using the fast and completely automated pipeline built around deep learning and Evolutionary Algorithms (EA) techniques^44^.

## Results and Discussion

Using the methods presented in^44^, we find 10 different sequences from the 10 runs of the EA technique. Next, we run an analysis to find if any of the sequences presents more than one mutation. The resulting sequence **GACCCACTTATGGT-GTTGGTC**presents 3 characteristics mutations to Omicron variant^1^: Q498R (A23055G), N501Y (A23063T) and Y505H (T23075C), position 23,054 to 23,075 in the reference accession NC_045512.2^45^. Then, we simulated sequence **GACC-CACTTATGGTGTTGGTC**as a forward primer in Primer3Plus^46^ using the accession *EPI*_*ISL*_6590782, resulting in a warning for *High end self complementarity*. To solve the issue, we increased the size of the primer by adding a bp at the end (**GACCCACTTATGGTGTTGGTCA**), which resulted in an acceptable primer candidate with a *T_m_* of 62.0 °C. We then generated the internal probe **CACCAGCAACTGTTTGTGGA**and reverse primer **CTGCCAAATTGTTGGAAAGG**with a *T_m_* of 60.8 °C and 60.5 °C respectively with a product size of 208 bp.

The frequency of appearance of the selected sequences is shown in Table 2. Out of 123 available Omicron sequences, 112 presented the candidate forward primer produced by our technique. From the 11 that do not present the sequence **GACCCACTTATGGTGTTGGTCA**, 9 had sequencing errors and 2 (1.63%) do not possess mutation Y505H. Further validation was performed on the different Omicron sub-lineages (BA.1, BA.2, BA.3) to verify that the primers work, Table 3. An analysis using 659 sequences, marked as *complete* and *low coverage excluded* in GISAID from BA.1 sub-lineage show 88.01% success of the forward primer and one nucleotide change in 2.28% of the sequences, while the rest have sequencing errors in the target.

**Table 2.**
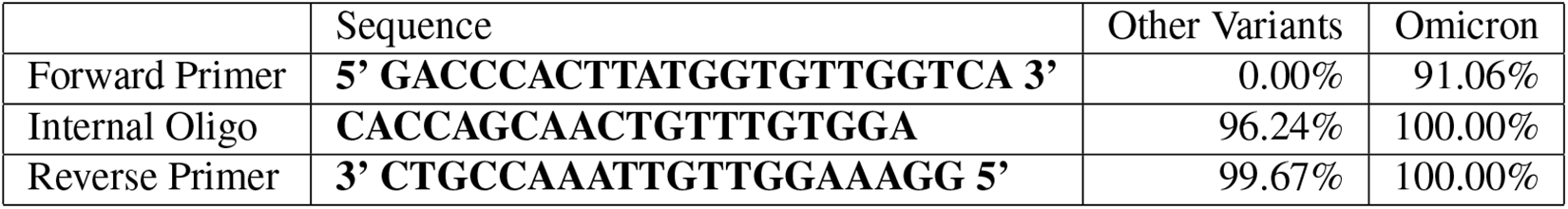
Frequency of appearance of the primer set in the sequences of the Omicron variant (123 samples) and other variants (2,100 samples).

**Table 3.**
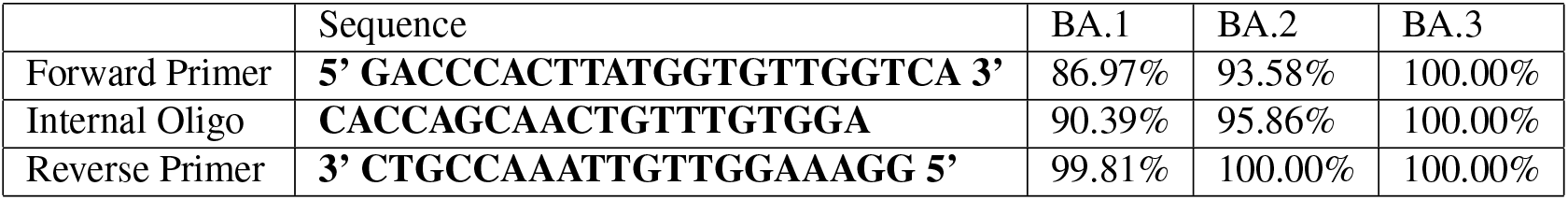
Frequency of appearance of the primer set in the sub-lineages of the Omicron variant: BA.1 (140,874 samples), BA.2 (701 samples), BA.3 (19 samples).

Laboratory testing of the generated primers, as part of the UniCoV study proves to be successful (see Table 4) in identifying the Omicron variant. Thus, these results prove further that the use of AI to generate specific diagnostic tests could be used as a fast measure to tackle the on-going SARS-CoV-2 pandemic and the appearance of new variants.

**Table 4.** qPCR results of the generated Omicron specific primers.

| Sample Code | N1 General Primer (CT) | Omicron Specific Primer (CT) | Omicron sample Y/N |
| --- | --- | --- | --- |
| 1515 | 33.96 | 37.00 | Y |
| 4702 | 28.79 | 33.04 | Y |
| 100308 | 36.06 | ND | N |
| 124546 | 26.61 | 32.37 | Y |
| 1122021a | 29.10 | ND | N |
| 105012022 | 28.69 | 36.51 | Y |
| Negative (dH2O) | ND | ND | N |
| RT-Negative | ND | ND | N |

## Methods and Data

### Data

From the Global Initiative on Sharing Avian Influenza Data (GISAID) repository^3^, we downloaded 123 sequences identified as B.1.1.529/Omicron and 100 sequences of each of the following variants labeled following the Pango lineage^2^, for a total of 2,100 sequences: AY.3 (Delta Sublineage), B.1, B.1.1.7 (Alpha), B.1.1.214, B.1.1.519, B.1.2, B.1.160, B.1.177, B.1.177.21, B.1.221, B.1.243, B.1.258, B.1.351 (Beta), B.1.427, B.1.429, B.1.526, B.1.596, B.1.617.2 (Delta), D.2, P.1 (Gamma), and R.1 in *.fasta format. Then we kept 23 sequences of B.1.1.529 as test set, while the other 100 were added to the 2,100 sequences of other variants to be used as a training set. The classification problem was defined as binary classification, with B.1.1.529 samples labeled with ‘1’, and the rest labeled with ‘0’. For further validation, we downloaded 140,874 sequences labeled as BA.1, 701 BA.2 and 19 BA.3 from the GISAID repository on January 6th, 2022, to verify whether our designed primer set works for the different Omicron sublineages.

### Methods

#### Creating the Primer Set

To create the primer set, we use a technique based on Evolutionary Algorithms (EAs), that is able to find suitable sequences to identify the Omicron variant^44^. In summary, this method optimizes two integer values *p, k* that jointly identify the best candidate forward primer of length 21 at position *p* in sample *k* of the training set, see Figure 1. Then gives a cost based on its frequency of appearance marked with label ‘1’, its absence from samples with label ‘0’ (in other words, its specificity to the target variant), its CG content, and melting temperature, as described in more detail in^47^, where the same technique was used to generate primers for B.1.1.7 (Alpha) variant successfully.

**Figure 1.**
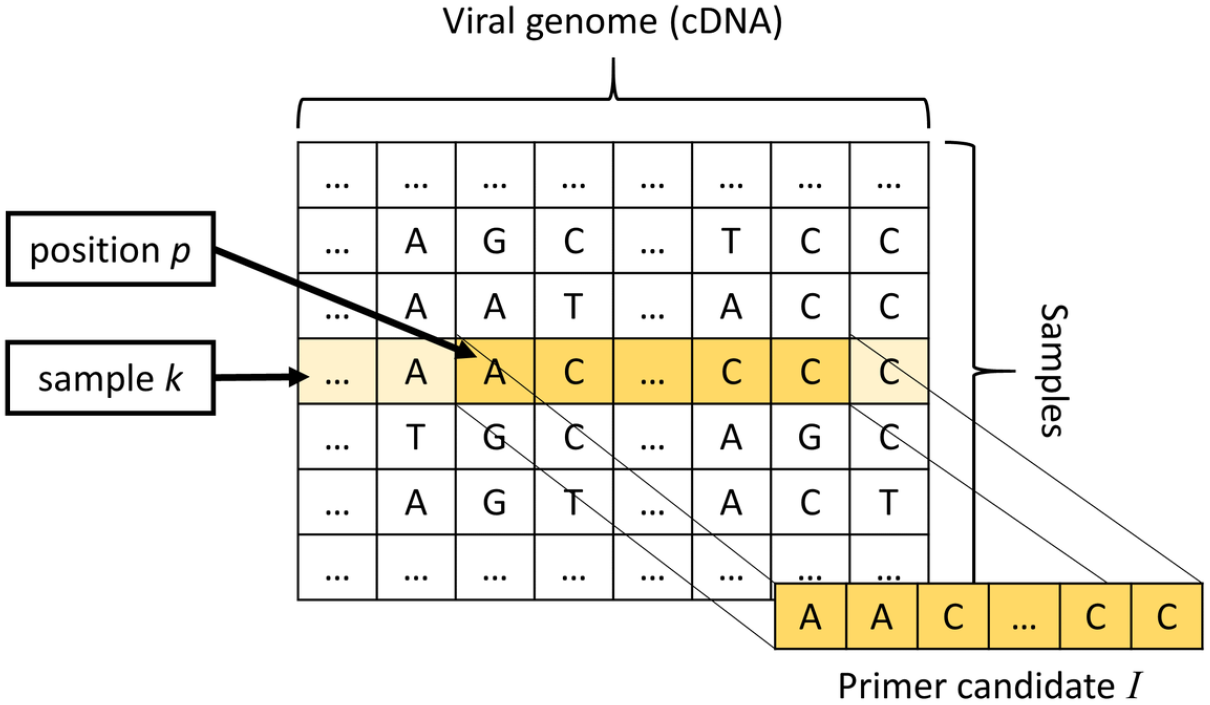
EA system where we select a target 21 bps sequence in position *p* and in sample *k*.

As EAs are stochastic algorithms, and can thus potentially return different candidates at each run, we run the algorithm 10 times to obtain a variety of candidate sequences. After the 10 repetitions, we check the obtained sequences for the number of mutations. Once a sequence containing more than one mutation is identified, we simulate it in Primer3Plus^46^ using the accession *EPI*_*ISL*_6590782 as a reference for Omicron variant.

#### Laboratory Testing

Raw Saliva samples, approx. 1-2 ml, were collected in 5 ml screwcap containers from volunteers in the UniCoV study^48^. Following collections samples were heat inactivated at 95°C for 5 min, cooled, vortexed and then 20 ml of Saliva was added to 20 ml of Saliva Ready™ Solution in a 0.2 ml 96 well plate. The plate was vortexed and centrifuged and then heated at 62°C for 5 min, 92°C for 5 min and then cooled at 4°C. Saliva was then screened for SARS-CoV-2 (ORF1a, ORF1b and N gene) and Human RNase P using the TaqMan™ 1-Step Mutiplex SARS-CoV-2 Fast PCR Kit 2.0 on the Applied Biosystems™ QuantStudio™ 5 Real Time PCR Instrument, 96 well, 0.2-mL block (Thermo Fisher Scientific) according to the manufacturer’s instructions including a Positive control. Reverse transcription 53°C for 5 min, 1 Preincubation 85°C for 5 min., Activation at 95°C 2 minutes with 40 cycles of Denaturation 95°C for 1 second and Anneal / extension 62°C for 30 seconds.

Positive SARS-CoV-2 samples were subsequently screened for the presence of Omicron variant using our specific Omicron Primer Set. Briefly cDNA was synthesised using saliva from the Saliva Ready™ step above with LunaScript RT-Supermix (NEB), briefly for primer annealing for 2 min at 25°C, cDNA synthesis for 10 minutes 55 °C 2 minutes and denaturation 95°C for 1 minute. qPCR using followed by second and Anneal / extension 62°C for 30 seconds.

qPCR was performed using cDNA from above with the Luna Universal Probe qPCR mix (NEB), with Omicron specific primers and N1 (2019-nCoV RUO) primers/probes (Integrated DNA Technologies (IDT)) at a concentration 500 nM with probes at 250 nM(FAM-labelled). PCR conditions were denaturation at 95°C fro 1 minutes and 40 cycles of Denaturation 95°C for 15 second and Anneal / extension 60°C for 30 seconds on the Applied Biosystems™ QuantStudio™ 5 Real Time PCR Instrument, 96 well, 0.2-mL block. All PCR reactions were performed in duplicate with a technical replicate performed following initial analysis.

The used data for this manuscript, and the resulting frequency of appearance for each sequence is available in: https://github.com/steppenwolf0/omicronVariant.git

## Author contributions statement

CAP, LMM, made the biological analysis, and primer design. ALR and AT made the programming, data collection, and experiments in silico. EC, ADK and JG made the experiment and study design. JMS and JS performed the wet lab validation on these primer sequences (from samples obtained as art of the UnCoV University community screening study in UCC, Cork, Ireland). CAP and ALR wrote the the article, all authors contributed to editing of the article.

## Additional information

J. Garssen is a part time employee at Danone Nutricia Research, Utrecht, the Netherlands.

## References

1. WHO. Classification of Omicron (B.1.1.529): SARS-CoV-2 Variant of Concern.

2. Rambaut, A. et al. A Dynamic Nomenclature Proposal for SARS-CoV-2 Lineages to Assist Genomic Epidemiology. Nat. microbiology 5, 1403–1407 (2020).

3. Shu, Y. & McCauley, J. GISAID: Global Initiative on Sharing All Influenza Data–From Vision to Reality. Eurosurveillance 22, 30494 (2017).

4. European Centre for Disease Prevention and Control. Implications of Emergence of the Emergence and Spread of the SARS-CoV-2 B.1.1.529 Variant of Concern (Omicron) for the EU/EEA.

5. Agency, U. H. S. SARS-CoV-1 variants of Concern and Variants under Investigation in England.

6. Tao, K. et al. The biological and clinical significance of emerging sars-cov-2 variants. Nat. Rev. Genet. 1–17 (2021).

7. Harvey, W. T. et al. Sars-cov-2 variants, spike mutations and immune escape. Nat. Rev. Microbiol. 19, 409–424 (2021).

8. Golcuk, M., Yildiz, A. & Gur, M. The omicron variant increases the interactions of sars-cov-2 spike glycoprotein with ace2. BioRxiv (2021).

9. Kumar, S., Thambiraja, T. S., Karuppanan, K. & Subramaniam, G. Omicron and delta variant of sars-cov-2: A comparative computational study of spike protein. J. medical virology (2021).

10. Lam, S. D., Waman, V. P., Orengo, C. & Lees, J. Insertions in the sars-cov-2 spike n-terminal domain may aid covid-19 transmission. bioRxiv (2021).

11. Omotuyi, O. I. et al. Sars-cov-2 omicron spike glycoprotein receptor binding domain exhibits super-binder ability with ace2 but not convalescent monoclonal antibody. bioRxiv (2021).

12. Cao, Y. R. et al. B. 1.1. 529 escapes the majority of sars-cov-2 neutralizing antibodies of diverse epitopes. bioRxiv (2021).

13. Brimacombe, K. R. et al. An opendata portal to share covid-19 drug repurposing data in real time. BioRxiv (2020).

14. Ford, C. T., Machado, D. J. & Janies, D. A. Predictions of the sars-cov-2 omicron variant (b. 1.1. 529) spike protein receptor-binding domain structure and neutralizing antibody interactions. bioRxiv (2021).

15. WHO. Update on omicron. https://www.who.int/news/item/28-11-2021-update-on-omicron.

16. Pulliam, J. R. et al. Increased risk of sars-cov-2 reinfection associated with emergence of the omicron variant in south africa. medRxiv DOI: 10.1101/2021.11.11.21266068 (2021).

17. CDC. Science brief: Omicron (b.1.1.529) variant.

18. Focosi, D., Franchini, M., Joyner, M. J. & Casadevall, A. Comparative analysis of antibody responses from covid-19 convalescents receiving various vaccines reveals consistent high neutralizing activity for sars-cov-2 variant of concern omicron. medRxiv (2021).

19. SARS-CoV-2 variants of concern and variants under investigation in England: Technical briefing 31 (2021).

20. COVID-19 Variant of Concern Omicron (B.1.1.529): Risk Assessment, December 7, 2021 (2021).

21. Covid, C. & Team, R. Sars-cov-2 b. 1.1. 529 (omicron) variant—united states, december 1–8, 2021. Morb. Mortal. Wkly. Rep. 70, 1731 (2021).

22. University, H. K. HKUMed finds Omicron SARS-CoV-2 can infect faster and better than Delta in human bronchus but with less severe infection in lung.

23. Discovery H. Discovery Health, South Africa’s largest private health insurance administrator, releases at-scale, real-world analysis of Omicron outbreak based on 211 000 COVID-19 test results in South Africa, including collaboration with the South Africa.

24. SARS-CoV-2 sequencing update: 1 December 2021 (2021).

25. Wolter, N. et al. Early assessment of the clinical severity of the sars-cov-2 omicron variant in south africa. medRxiv (2021).

26. Carreno, J. M. et al. Activity of convalescent and vaccine serum against a b. 1.1. 529 variant sars-cov-2 isolate. medRxiv (2021).

27. SARS-CoV-2 variants of concern and variants under investigation in England: Technical briefing 33 (2021).

28. Liu, L. et al. Striking antibody evasion manifested by the omicron variant of sars-cov-2. bioRxiv (2021).

29. Stumpp, M. Ensovibep, a novel trispecific darpin candidate that protects against sars-cov-2 variants. bioArxiv (2021).

30. Vangeel, L. et al. Remdesivir, molnupiravir and nirmatrelvir remain active against sars-cov-2 omicron and other variants of concern. bioRxiv DOI: 10.1101/2021.12.27.474275 (2021).

31. Lusvarghi, S. et al. Sars-cov-2 omicron neutralization by therapeutic antibodies, convalescent sera, and post-mrna vaccine booster. bioRxiv DOI: 10.1101/2021.12.22.473880 (2021). https://www.biorxiv.org/content/early/2021/12/28/2021.12.22.473880.full.pdf.

32. Andrews, N. et al. Effectiveness of covid-19 vaccines against the omicron (b. 1.1. 529) variant of concern. medRxiv (2021).

33. Edara, V.-V. et al. mrna-1273 and bnt162b2 mrna vaccines have reduced neutralizing activity against the sars-cov-2 omicron variant. bioRxiv (2021).

34. Wang, X. et al. Homologous or heterologous booster of inactivated vaccine reduces sars-cov-2 omicron variant escape from neutralizing antibodies. bioRxiv DOI: 10.1101/2021.12.24.474138 (2021). https://www.biorxiv.org/content/early/2021/12/27/2021.12.24.474138.full.pdf.

35. GeurtsvanKessel, C. H. et al. Divergent sars cov-2 omicron-specific t- and b-cell responses in covid-19 vaccine recipients. medRxiv DOI: 10.1101/2021.12.27.21268416 (2021). https://www.medrxiv.org/content/early/2021/12/29/2021.12.27.21268416.full.pdf.

36. Garcia-Beltran, W. F. et al. mrna-based covid-19 vaccine boosters induce neutralizing immunity against sars-cov-2 omicron variant. Cell (2021).

37. Dolzhikova, I. V. et al. Sputnik light booster after sputnik v vaccination induces robust neutralizing antibody response to b. 1.1. 529 (omicron) sars-cov-2 variant. medRxiv (2021).

38. Malone, B. et al. Artificial intelligence predicts the immunogenic landscape of sars-cov-2 leading to universal blueprints for vaccine designs. Sci. reports 10, 1–14 (2020).

39. Khan, M. T. et al. Immunoinformatics and molecular modeling approach to design universal multi-epitope vaccine for sars-cov-2. Informatics medicine unlocked 24, 100578 (2021).

40. Khan, K. et al. Omicron infection enhances neutralizing immunity against the delta variant. medRxiv DOI: 10.1101/2021.12.27.21268439 (2021).

41. Janse, M., Brouwers, T., Claassen, E., Hermans, P. & Van De Burgwal, L. Barriers influencing vaccine development timelines, identification, causal analysis, and prioritization of key barriers by kols in general and covid-19 vaccine r&d. Front. public health 9, 395 (2021).

42. van der Waal, M. B. et al. Blockchain-facilitated sharing to advance outbreak r&d. Science 368, 719–721 (2020).

43. Chand, M., Hopkins, S. & Dabrera, G. Investigation of novel SARS-COV-2 variant: Variant of Concern 202012/01 (2020).

44. Lopez-Rincon, A. et al. Design of Specific Primer Sets for the Detection of B.1.1.7, B.1.351, P. 1, B. 1.617. 2 and B. 1.1. 519 Variants of SARS-CoV-2 using Artificial Intelligence. bioRxiv (2021).

45. Wang, C. et al. The Establishment of Reference Sequence for SARS-CoV-2 and Variation Analysis. J. medical virology 92, 667–674 (2020).

46. Untergasser, A. et al. Primer3Plus, an Enhanced Web Interface to Primer3. Nucleic acids research 35, W71–W74 (2007).

47. Rincon, A. L. et al. Design of Specific Primer Sets for SARS-CoV-2 Variants Using Evolutionary Algorithms. In Proceedings of the Genetic and Evolutionary Computation Conference, 982–990 (2021).

48. UCC unicov. https://www.ucc.ie/en/emt/covid19/uni-cov/.

